# Rational Design of Evolutionarily Stable Microbial Kill Switches

**DOI:** 10.1101/129445

**Authors:** Finn Stirling, Lisa Bitzan, John WK Oliver, Jeffrey Way, Pamela A. Silver

**Affiliations:** Department of Systems Biology, Harvard Medical School, 200 Longwood Avenue, Warren Alpert 536, Boston, MA 02115, USA; Wyss Institute for Biologically Inspired Engineering, Harvard University, 3 Blackfan Circle, 5th Floor, Boston, MA 02115, USA

**Keywords:** Containment, Library, Synthetic Biology, Lambda, CspA, Promoter, Toxin, Antitoxin, Cold Shock

## Abstract

The evolutionary stability of synthetic genetic circuits is key to both the understanding and application of genetic control elements. One particularly useful but challenging situation is a switch between life and death depending on environment. Here are presented “essentializer” and “cryodeath” circuits, which act as kill switches in *Escherichia coli*. The essentializer element induces cell death upon the loss of a bi-stable cI/Cro memory switch. Cryodeath makes use of a cold-inducible promoter to express a toxin. We employ rational design and a novel toxin/antitoxin titering approach to produce and screen a small library of potential constructs, in order to select for constructs that are evolutionarily stable. Both kill switches were shown to maintain functionality in vitro for at least 140 generations. In addition, cryodeath was shown to control the growth environment of a bacterial population, with an escape rate of less than 1 in 10^5^ after ten days *in vivo*.

## Introduction

As synthetic biology makes advances in producing real world applications using genetically engineered micro-organisms, the issue of biological containment becomes increasingly important. Safeguards have previously been developed that require the addition of a survival factor to maintain viability in a bacterial population. Approaches include inducing an auxotrophy for a particular metabolite (Gallagher, Patel, Interiano, Rovner, & Isaacs, 2015; L Steidler et al., 2003; Wright, Delmans, Stan, & Ellis, 2015), repressing the expression of an essential gene (Cai et al., 2015; Chan, Lee, Cameron, Bashor, & Collins, 2015; Gallagher et al., 2015) or re-writing the genetic code to ensure dependency on an artificial amino acid (Isaacs et al., 2011). However, future applications of synthetic biology look beyond the confines of a laboratory. In research settings, strains have already been developed that can degrade inorganic polymers to reduce waste (Yoshida et al., 2016), provide sustenance or energy during space travel (Menezes, Cumbers, Hogan, & Arkin, 2014; Montague et al., 2012; Way, Silver, & Howard, 2011), or that colonize the mammalian gut to help diagnose and treat pathogenic infections (Kotula et al., 2014; Lothar Steidler, 2003). For these applications, a new form of containment is required for uncontrolled environments, one that does not require human monitoring or input.

Evolutionary instability is an inherent flaw in any biological control using a lethal effect to control environmental growth. Microorganisms rapidly evolve to remove any genetic element that reduces fitness (Knudsen & Karlstrom, 1991; Molin et al., 1993). “Kill switches” are defined as artificial systems that result in cell death under certain conditions. Several kill switches have been explored for containment of engineered microbes, but necessarily involve lethal genes that are induced in designated non-permissive conditions. Thus any kill switch with leaky, low level expression of a toxin in supposedly permissive conditions may be quickly disabled in rapidly growing microbes.

Although a host of highly lethal and effective kill switches have been described, most evolve to lose functionality within days (Chan et al., 2015), or have no data supporting their longevity (Callura, Dwyer, Isaacs, Cantor, & Collins, 2010; Piraner, Abedi, Moser, Lee-Gosselin, & Shapiro, 2016). One notable exception is Gallagher et al’s publication on multi layered kill switches (Gallagher et al., 2015), which details a system stable for at least 110 generations. However, this system relies upon being able to provide the required survival factors to the engineered strain.

To create evolutionarily stable kill switches, we use a novel strategy for varying the level of expression in a toxin/antitoxin system. Small rationally designed libraries were created with key bases modified in the promoter and RBS sites of both toxin and antitoxin. This system is broadly applicable in kill switch designs with diverse forms of regulation. We demonstrate this approach with two unrelated control systems: (1) the early termination of transgenics that lose function of another engineered module; and (2), the temperature-dependent termination of a transgenic microorganism that resides in the mammalian gut. In doing so, we explore a limited rationally designed space to achieve the optimized behavior of genetic circuits.

## Results

### General design strategy

An environmentally sensitive kill switch needs to be highly lethal in non-permissive conditions, but stable enough in permissive conditions to avoid conferring an evolutionary disadvantage. Achieving intended levels of protein repression can be more straightforward when a molecular ‘sponge’ is included to titrate low levels of an expressed toxin (Bläsi & Young, 1996; Nathan, Glenn, & Johan, 2016). In our system, this has been achieved through toxin/antitoxin pairings where the antitoxin sequesters toxin expressed during permissive states, but does not prevent cell death after a non-permissive event (Figure 1A). The toxin is placed under the control of an environmentally sensitive promoter, capable of strong repression in permissive conditions and high levels of expression in non-permissive conditions (Figure 1B). Even with highly stringent repression of a promoter, there will always be leaky expression of the intended gene product. For a toxin lethal enough to make an effective kill switch, any unintended expression would have a negative effect on cell survival and decrease evolutionary stability.

**Figure 1:**
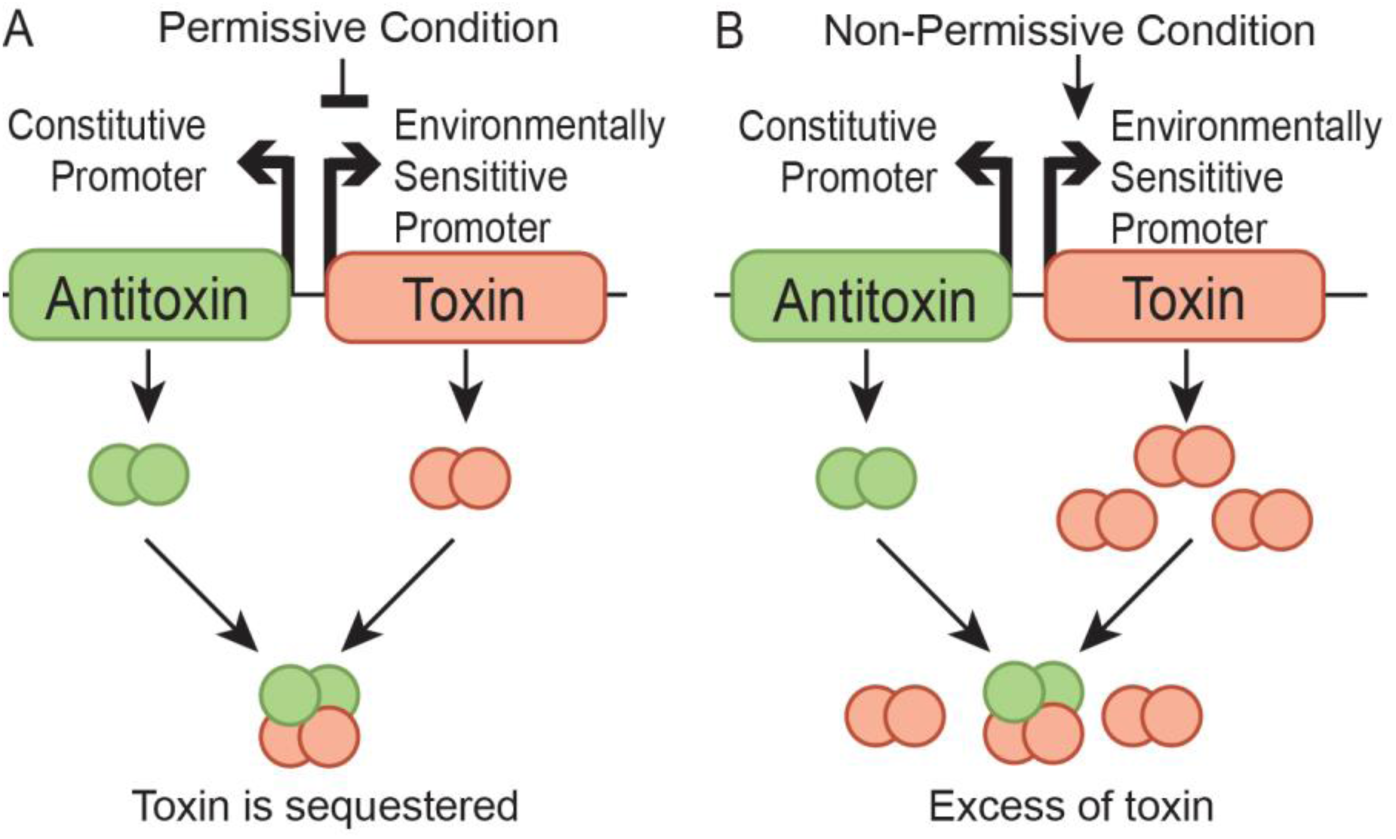
Overview of kill switch design concept. **A)** In permissive conditions the toxin is repressed. The antitoxin is expressed at a constitutive low level to accommodate for any leaky expression of the toxin. **B)** Upon a change of environment to non-permissive conditions the repression is lifted and toxin expression increases. The low level of antitoxin expression is no longer capable of preventing a lethal level of free toxin.

The toxin-antitoxin system ccdB/ccdA allows for repression of toxic effects in permissive conditions with a high cell death rate in non-permissive conditions. CcdB is a lethal toxin for many Enterobacteriaceae (Wright, Stan, & Ellis, 2013). It targets the GyrA subunit of DNA gyrase, arresting it in the intermediate stage of action after a double-stranded break has been induced (Madl et al., 2006). With no mechanism for repairing a double-stranded break, cell death results (Bahassi, O’Dea, Allali, Messens, & Gellert, 1999). CcdB/CcdA is a well characterized system, found natively as a suicide factor on the F plasmid in *E. coli* (Bernard & Couturier, 1992).

### An ‘Essentializer’ Element to Select for the Preservation of a Transgenic Cassette

In previous work, we have published a bistable genetic switch, the ‘memory element’, that is able to stably record and maintain an output from an exogenous signal (Kotula et al., 2014). The memory element is required to maintain its function for extensive periods of time, over which synthetic circuits are prone to mutation and deletion (Sleight, Bartley, Lieviant, & Sauro, 2010). The memory element relies upon the bacteriophage lambda transcription factors cI and Cro. cI binds preferentially to the operator regions O_R_1 and O_R_2, whilst Cro binds preferentially to O_R_3. In lambda and in the memory element, binding of cI to O_R_1 and O_R_2 represses the expression of *cro*, and vice versa binding of Cro to O_R_3 represses the expression of *cI*.

The essentializer element kill switch was designed using the toxin-antitoxin system CcdB/CcdA, and would select for the presence of the memory element. For the essentializer element, the lambda phage operator sites were reordered and positioned over the lambda P_R_ promoter such that binding of either cI or Cro would repress the expression of the toxin (Figure 2A and 2B). Furthermore, the binding sites O_R_1 and O_R_2 were modified to increase the binding affinity of cI (Sarai & Takeda, 1989; Takeda, Sarai, & Rivera, 1989). A modified O_R_3 was placed downstream of the -10 region for the P_R_/toxin promoter in a position analogous to *lacO* within *lacP*; a protein binding to the major groove of these bases is expected to sterically prevent RNA polymerase binding (Murakami, Masuda, Campbell, Muzzin, & Darst, 2002). The operators are separated by 6-7 bases, which should allow cooperative binding of cI to either O_R_1-O_R_2 or O_R_1-O_R_3 in this configuration (Ptashne et al., 1980).

**Figure 2:**
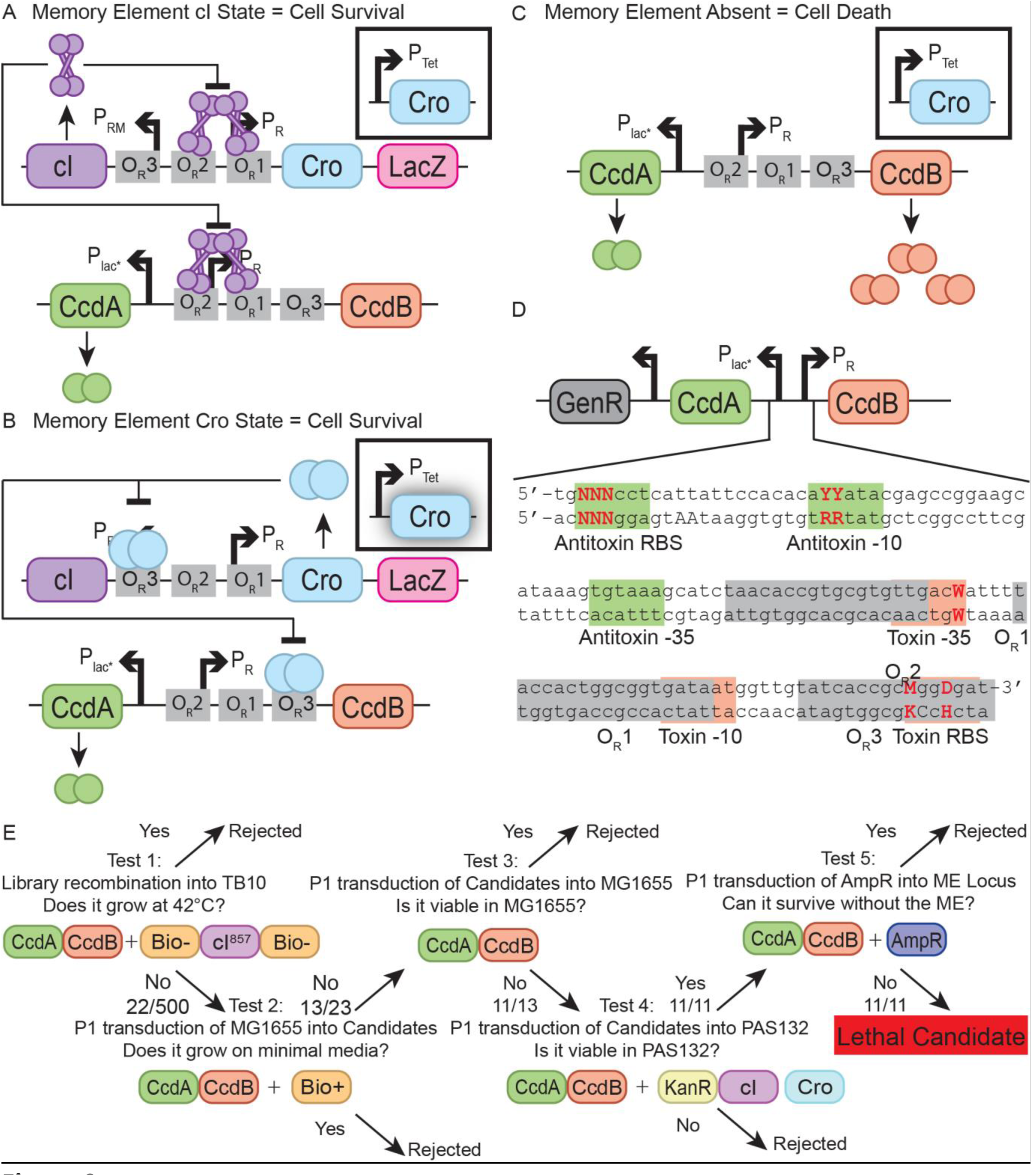
Design and construction of the essentializer element kill switch. **A)** Memory element (top cassette) in the “off” configuration. Expression of *cI* represses expression of *cro* and *lacZ*, whilst simultaneously repressing expression of *ccdB* in the essentializer element (bottom cassette). *ccdA* is expressed at a low constitutive level to accommodate for any leaky expression from the toxin. Additionally *cro* is expressed under a tetracycline sensitive promoter that acts as a trigger to switch between the *cI* and *cro* states of the memory element. **B)** Exposure to tetracycline leads to a pulse of expression of *cro* from the trigger element, setting the memory element in the “on” configuration. Expression of *cro* allows for the expression of *lacZ*, whilst simultaneously repressing *cI* and *ccdB*. **C)** Memory element is absent. Without repression from *cI* or *cro*, *ccdB* is expressed at lethal levels. **D)** The engineered region encompassing the regulatory DNA for *ccdB* and *ccdA*. Highlighted are loci that have a strong impact on expression level for both the toxin (red) and antitoxin (green), as well as operator binding sites for cI and Cro (grey). Key bases that have been varied are bolded in red and capitalized. By varying these bases, a library of constructs with differing levels of expression was created and subsequently screened to identify the construct with the desired phenotype: N = A, C, T or G; Y = C or T, W = A or T; M = A or C; D = A, T or G. **E)** Overview of the screening process for identifying lethal essentializer element candidates. Each step conducted a different test to select for candidates that had potential to be a lethal essentializer element. The relevant genotype after each step has been completed is displayed, along with the number of candidates that passed a given screen/number of candidates that were tested.

In the memory element’s dormant *cI* state (Figure 2A), cI is expressed, repressing both *cro* and *ccdB*, which should allow for cell survival. When the memory element is switched to its active Cro state by means of a trigger element (Figure 2B), *cro* is expressed by the memory element, repressing both *cI* and *ccdB*, and also allowing for cell survival and the expression of a reporter gene by the memory element. In particular, binding of Cro to O_R_2, O_R_1 or O_R_3 should be sufficient to repress toxin synthesis. However, if the memory element is lost entirely from the cell (Figure 2C) neither *cI* nor *cro* will be expressed, allowing for expression of *ccdB* and cell death. The antitoxin CcdA is expressed under a constitutive promoter based on the LacUV5 promoter (Plac*), which was chosen because it is roughly as strong as the lambda P_R_ (Simons, Hoopes, McClure, & Kleckner, 1983), to accommodate for any leaky expression of the toxin under permissive conditions.

A specific design goal was to quantitatively adjust the levels of toxin and antitoxin expression so that when cI and Cro proteins are absent, toxin expression is sufficient to kill the cell, but when either Cro or cI protein is present, the toxin is sufficiently repressed such that cell growth is not affected. This should be true even allowing for stochastic binding of repressor or Cro to the operators, and stochastic expression of the antitoxin. To achieve this goal, a small rationally designed library of essentializer element candidates was constructed to introduce degeneracy at key locations in the regulatory region of the kill switch (Figure 2D). Three bases were varied in the RBS for the antitoxin, each with 4 different possible nucleotides; two bases were varied in the -10 region of the antitoxin promoter, each with 2 different possible nucleotides; one base was varied in the -35 region of the toxin promoter, with 2 different possible nucleotides; two bases were varied in the RBS for the toxin, one with 2 possible nucleotides and the other with 3. This resulted in a library size of 4^3 × 3^1 × 2^4 = 3072 potential combinations. The promoter variations were designed based on Mulligan et al. (Mulligan, Hawley, Entriken, & McClure, 1984), who estimated that, for example, the possible non-preferred bases in the 4^th^ and 5^th^ positions in the consensus -10 region (TAT**<u>AA</u>**T) cause a mild and roughly equal decrement in promoter strength, so that incorporating only two variants would generate maximal functional variation while limiting the number of candidates to be manually screened. Variations in the toxin RBS were chosen to potentially preserve cI and Cro-binding at the overlapping O_R_3 element (Sarai & Takeda, 1989; Takeda et al., 1989). A construct was also made with a frame shift mutation in the toxin open reading frame to act as a non-lethal control (“EE toxin mutant”).

The essentializer element candidate pool yielded constructs that depended upon the memory element for survival. A series of tests was designed to select for candidates that survived in the permissive conditions (presence of memory element in the *cI* state) but failed to grow in non-permissive conditions (absence of memory element) (Figure 2E). In test 1, the essentializer element library was recombined into the lambda *red* expressing strain TB10. TB10 expresses *cI*^857^, a temperature sensitive mutant of *cI* (Caulcott & Rhodes, 1986) which is active at 30 °C but inactive at 42 °C, therefore repressing *ccdB* in a temperature dependent manner. Any recombinant that failed to grow at 42°C but survived at 30°C was considered potentially lethal. Test 2 used a P1 transduction of a wild type MG1655 lysate to remove *cI*^857^, allowing the strain to regain an operational biotin operon. Successful growth on minimal media without biotin therefore implied a non-lethal essentializer element as cI^857^ would no longer be present to repress. Test 3 involved P1 transducing each essentializer element candidate into MG1655, relying on the distance between the loci of *cI*^857^ and the essentializer element to prevent them being cotransduced. With no source of cI or Cro in MG1655, if a candidate failed to produce any transductants, it was a potentially viable candidate. Test 4 transduced the essentializer elements into PAS132, a strain containing the memory element (Kotula et al., 2014) preset to the cI state. If a candidate produced transductants here, but had failed to produce transductants in test 3, it was a potentially viable candidate. Candidates were sequenced across the antitoxin, toxin and regulatory region to determine the exact sequence (Supplementary Figure 1), and termed candidates EE01-EE11. All other sequenced candidates were termed EE12-EE31.

The new candidate strains containing the essentializer element in a PAS132 background showed selection for maintaining the memory element. The fifth test inserted ampicillin resistance by P1 transduction into the candidate strains approximately 3000 bp distal to the memory element, likely replacing it. The more common transductant of removing the memory element would result in only ampicillin resistance and the essentializer element remaining, but a rare transduction event could insert ampicillin resistance without removing the memory element and its associated kanamycin resistance cassette (Figure 3A). If a candidate contained a functional essentializer element, it would not be able to survive without the memory element. So only the rare transductant would be viable, whereas controls would have a majority of common transductants with a low proportion of rare transductants. In all 11 candidates, only the rare transductant result was recovered out of 52 transductants screened for growth on ampicillin and subsequently restreaked onto kanamycin (Figure 3B red bars).

**Figure 3:**
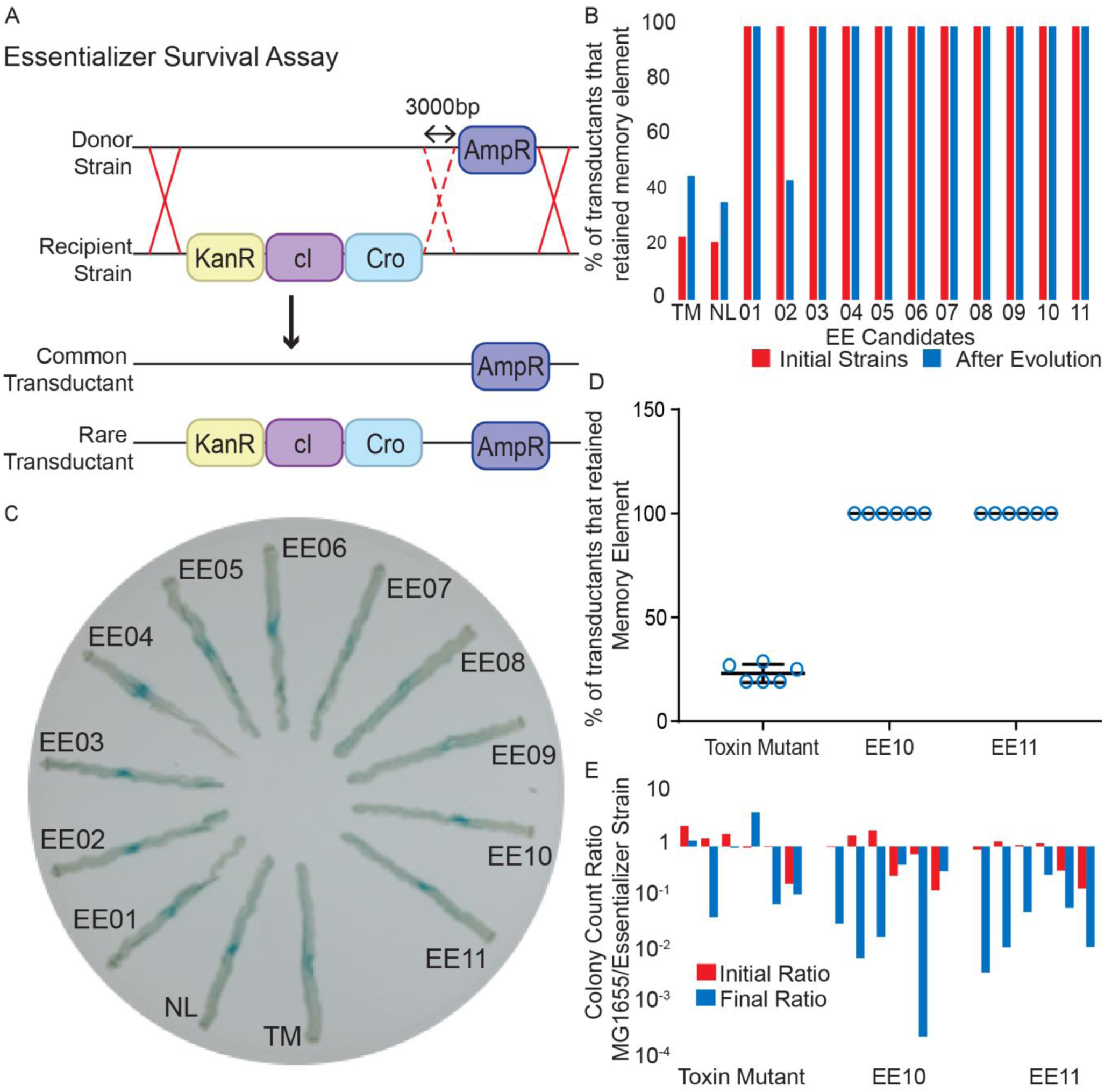
Analysis of essentializer element candidates: **A)** Diagram detailing the removal of the memory element. A P1 transduction donor strain with ampicillin resistance ∼3000 bp from the memory element was used to remove the memory element. Due to the spacing between the loci of the cassettes a small subset of transductants would have both cassettes. **B)** Percentage of colonies that retained the memory element after the assay outlined in figure 6A, both before (red) and after (blue) passaging for 140 generations. TM = toxin mutant and NL = EE non-lethal. **C)** Candidate essentializer strains were streaked on an agar plate in which a spot of ATc was placed at the center, resulting in a diffusion gradient. The result is to induce high-level Cro expression near the center and repress *lacZ* expression from the memory element (Figure 2A,B), to induce switching to a stable Cro state within the memory element and allow *lacZ* expression at an intermediate ATc level, and to remain in the cI state at a low ATc level. Lethality due to the essentializer is not observed in any of the Cro or cI expression states. **D)** Six biological repeats of the assay in 6A for candidates EE10 and EE11. **E)** Six biological repeats of a competitive growth assay to compare the fitness of parental (MG1655) and engineered bacterial strains. Mixed cultures were titered before and after sub-culturing for 70 generations.

The lethal activity of essentializer element candidates remained stable over extended growth periods in the cI state (Figure 3B). Candidates were passaged for approximately 140 generations and then the test outlined in Figure 3A was used to assay whether the memory element was still essential. Only candidate EE02 failed to maintain its selection for preservation of the memory element (Figure 3B); subsequent sequencing revealed multiple mutations in the regulatory region of both the toxin and the antitoxin for candidates EE02 and EE06.

Upon switching to the Cro state of the memory element, candidates maintained viability with the presence of Cro rather than cI. PAS132 contains a trigger element, allowing for the expression of *cro* under a tetracycline sensitive promoter P_Tet_ (Kotula et al., 2014). If grown in the presence of anhydrotetracycline (ATc), the memory element switches from the cI state to the Cro state through expression of the P_*Tet*_*cro* trigger. All candidates were streaked on a gradient of ATc spanning sub-induction to super-induction levels. At high ATc concentrations, Cro binds to the O_R_1 and O_R_2 operator regions in addition to its preferred O_R_3, repressing itself in the memory element design. For this reason, switching to the Cro state can only occur in a narrow range of Cro concentrations. In Figure 3E we see narrow strips of blue for each candidate, indicating the strain has survived the memory element flipping from the cI state to the Cro state, and subsequently allowed expression of the reporter beta-galactosidase (LacZ).

The essentializer element candidates EE10 and EE11 were selected for more in depth assays of their evolutionary stability. Each candidate was passaged for approximately 140 generations, along with the toxin-defective control. The candidates were subsequently tested for transductional removal of the memory element as outlined in Figure 3A. For both EE10 and EE11, 52 of 52 colonies screened for each biological repeat maintained the memory element, compared with about 20% for the toxin mutant control (Figure 3C). In addition, co-cultures of roughly equal starting ratio of the essentializer candidates and the parental MG1655 strain were grown and passaged for approximately 70 generations. The ratio of essentializer strain to parental MG1655 was measured by comparing colony-forming units before and after growth across 6 biological repeats (Figure 3D). Strains EE10 and EE11 as well as the toxin mutant generally had a greater number of cells present than MG1655, implying that the essentializer element does not confer a significant growth disadvantage.

The sequences of the artificial regulatory regions were determined in essentializer elements EE01-EE11, as well as for several elements that failed at various points during the screening process. Overall, about 2/3 of the sequences had deletions of 1 or more bases in this region. The remainder of *ccdA* and *ccdB* were not sequenced, and it is possible that additional mutations could have occurred in these genes or in the host in some cases. In five cases (the lethal candidates EE01 and EE07 as well as the 4 non-lethal candidates EE13, EE15 and EE27) the 5’-most operator (O_R_2) has a deletion and yet the strains are always or conditionally viable, indicating that loss of this operator is not required for tight repression and suggesting that the system is somewhat overdesigned. Beyond this, it is difficult to define a structure-activity relationship between the regulatory region sequences and the phenotypes observed, possibly because the widely varying ribosome binding sites may lead to changes in mRNA structure that are hard to predict.

### Design and Construction of the Temperature Sensitive Kill Switch ‘Cryodeath’

To further test our approach to balancing toxin expression, we designed a kill switch that would respond to temperature. The temperature sensitive regulatory region of cold shock protein A (P_*cspA*_) is modular and confers temperature-regulated expression of a protein of interest (Lee et al., 1994). P_*cspA*_ contains a constitutive promoter that has a high rate of transcription at all temperatures (Yamanaka, 1999). This is followed by a long 5’ untranslated region (UTR) of 159 bp that adopts an unstable secondary structure at 37°C and is rapidly degraded by RNaseE (Mitta, Fang, & Inouye, 1997), but at lower temperatures forms a stable configuration that allows translation (Fang, Jiang, Bae, & Inouye, 1997; Giuliodori et al., 2010). The cold-shock regulatory region also includes a downstream box (DB) element located within the first 13 amino acids of the open reading frame of *cspA* that enhances translation during cold shock by binding to the anti-DB sequence of 16S rRNA (Etchegaray & Inouye, 1999; Mitta et al., 1997) (Figure 4A).

**Figure 4:**
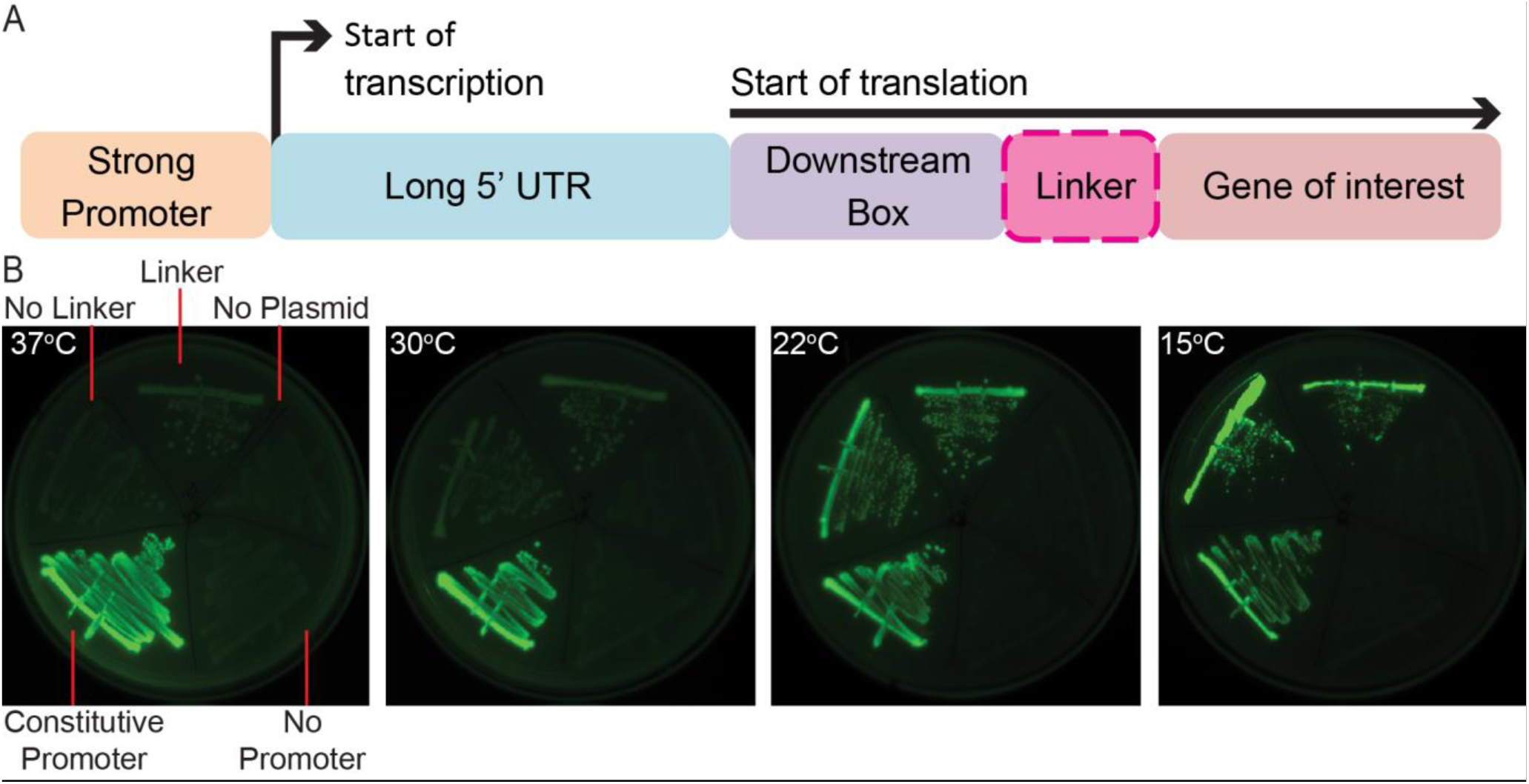
Structure and modularity of P_*cspA*_. **A)** The regulatory region of cold shock protein A (CspA). A strong promoter leads to a high constitutive level of transcription at all temperatures. The long 5’ UTR adopts an unstable configuration at high temperatures and is quickly degraded. At low temperatures the downstream box within the open reading frame of *cspA* assists in the binding of the 16S ribosomal subunit. This entire regulatory region can be placed upstream of a gene of interest to make its expression temperature sensitive, and connected either with or without a linker to avoid potentially adverse effects of forming a chimeric protein. **B)** Expression of GFP under P_*cspA*_ at 37^°^C, 30^°^C, 22^°^C, and 15^°^C for 10 days. GFP was expressed on the plasmid vector pKD46 under P_*cspA*_ either with or without a linker and compared to three controls, expression under the constitutive promoter P_*rpsL*_, No Promoter and no Plasmid. Images are from the same picture, taken on a Biorad ChemiDoc XRS+.

We demonstrated the utility of this isolated regulatory region by inducing GFP in a temperature sensitive fashion. As it was unclear whether a particular gene of interest would be affected by an additional 13 amino acids on the amino end, we generated two versions of the regulatory region fused to GFP - one with a linker (N-GGGGS-C) between the truncated CspA and GFP designed to minimize interaction of the DB element, and one with no linker (Figure 4A). When assayed for expression at 37°C, no discernible expression was observed compared to negative controls. However, to a small extent at 30°C, and greater extent at room temperature (referred to as 22°C) and 15°C, the level of induction was visibly increased both with and without the linker (Figure 4B), suggesting that the regulatory region functioned in a temperature sensitive manner when used to express GFP.

Rational design was used to generate small libraries to introduce variation at key locations in the regulatory region of the toxin and antitoxin. The *ccdB* coding region was placed after the *cspA* regulatory region, both with and without a linker. In a similar design to the essentializer element, the *ccdA* coding region was placed after a modified, constitutive LacUV5 promoter (P_*lac*_*) (Malan & McClure, 1984) (Figures 5A and 5B). A Gentamycin resistance cassette was used for selection purposes. To achieve a range of expression levels for the toxin and antitoxin, ten bases were varied - three in the RBS of the antitoxin, two in the -10 promoter region of the antitoxin, one in the -35 promoter region of the toxin, two in the -10 promoter region of the toxin, and two in the RBS of the toxin (Figure 5C). Each varied position had the possibility of 2 different nucleotides, making a possible 2^10 = 1024 different constructs. The bases chosen to be varied followed a similar logic to that of the essentializer element, generally with bases of less importance from within the -10 and -35 regions, according to previous published analysis of *E. coli* promoter and ribosome binding sites (Mulligan et al., 1984; R K Shultzaberger, Bucheimer, Rudd, & Schneider, 2001; Ryan K. Shultzaberger, Chen, Lewis, & Schneider, 2007).

**Figure 5:**
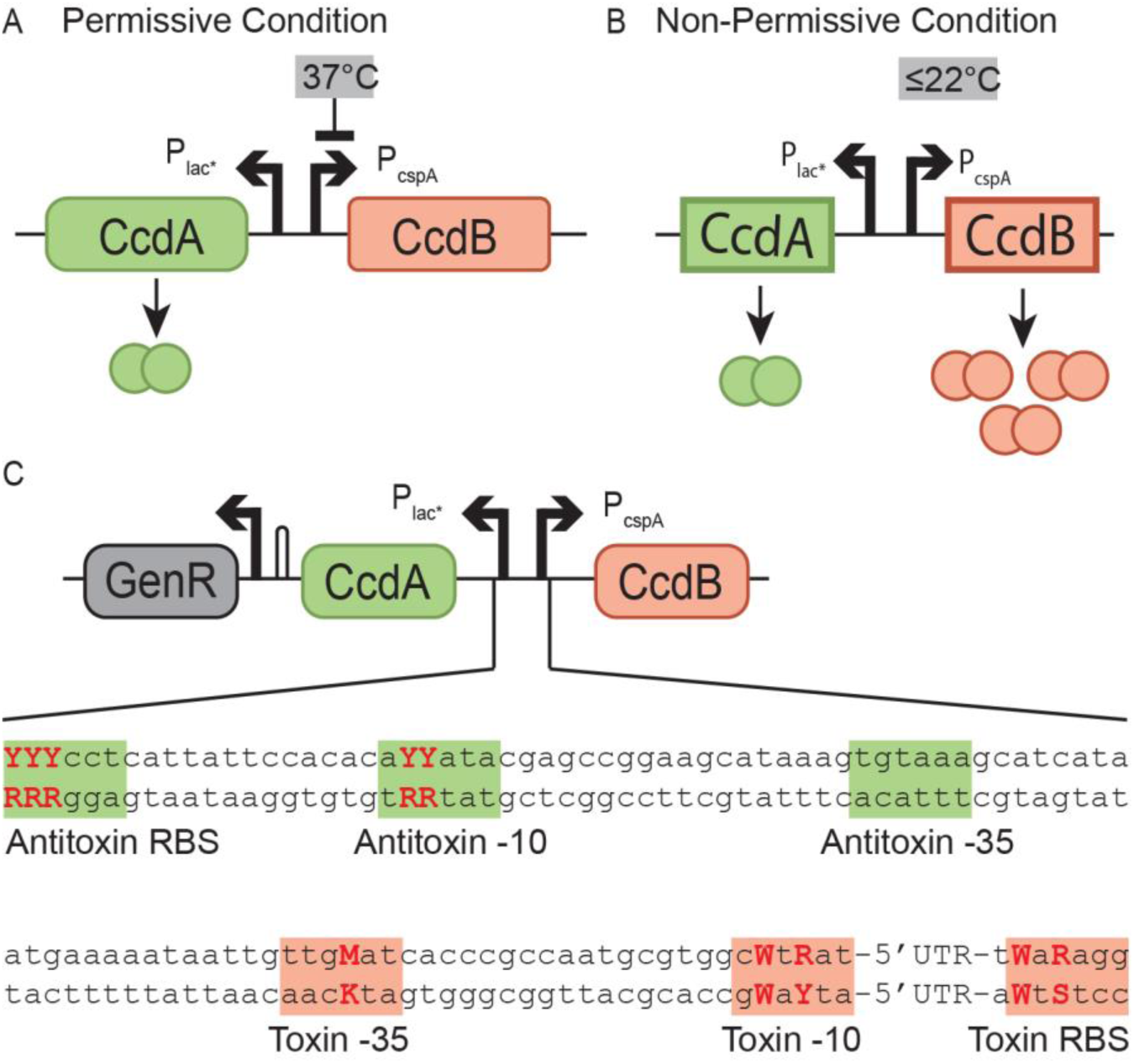
Design of the temperature sensitive kill switch cryodeath. **A)** At 37 °C the mRNA of the toxin, CcdB, is rapidly degraded due to an unstable 5’ UTR secondary structure, limiting translation and allowing cell survival. **B)** At colder temperatures, the 5’ UTR adopts a more stable conformation allowing translation of CcdB. **C)** The engineered region encompassing the regulatory DNA for CcdB and CcdA. Highlighted are segments that have a strong impact on expression level for both the toxin (red) and antitoxin (green). Varied bases are bolded in red and capitalized. By varying these bases, a library of constructs with differing levels of expression was created and subsequently screened to identify the construct with the desired phenotype. Y = C or T; M = A or C; W = A or T; R = A or G.

To screen for cold-sensitive toxin induced death, *E. coli* strain Dh10β was transformed with the linear fragment libraries, using the lambda *red* genes expressed on plasmid pKD46 to enhance recombination. After an initial selection for gentamicin resistance, 26 unique candidates were identified; 2 with no linker and 24 with a linker. The 26 candidates were colony-purified on LB agar and tested on plates at the non-permissive temperatures of 30°C, 22^o^ and 15°C as well as the permissive temperature, 37°C. Of the 26 candidates, ten failed to grow or grew poorly at <37°C, and were selected for further analysis (CD01-CD10). Non-lethal candidates were termed CD11-CD26. All except CD01 and CD11 contained a linker. As lethal candidates were identified btoh with and without a linker, it is probably that the linker has no or little effect on the function of CcdB. CD12 was subsequently used as a negative control, referred to in this text as “CD non-lethal”. The candidates were sequenced across the antitoxin, toxin, and regulatory region, toxin and antitoxin open reading frames (supplementary table 6). Although most candidates had retained the expected sequence across the regulatory region, only varying at the intended positions, candidate CD10 had a 9bp deletion partially including and upstream of the toxin RBS (supplementary figure 3). Modifying bases in this region has previously been noted to affect the stability of the CspA mRNA (Fang et al., 1997). In addition, CD01 was missing the third and fourth codons of the antitoxin (supplementary figure 3).

Evolutionarily stable circuits with the intended phenotype were identified. A survival assay comparing cfu at room temperature to that at 37°C was used to quantify the extent of population death, termed the survival ratio (Figure 6A). After an initial assay was made at room temperature, the candidates were passaged in permissive conditions for approximately 140 generations in LB before a second assay for cold-sensitivity was performed at room temperature. The pre- and post-survival ratios for each candidate were compared to determine which candidates induced the highest level of population death, and which candidates maintained their lethality after the opportunity for evolution (Figure 6B). CD01 and CD10 had low survival ratios of approximately 10^-4^ both before and after the period of growth, and were therefore chosen for further analysis. All candidates were sequenced across the antitoxin, toxin and regulatory region after the period of growth, with no mutations. For the candidates which appear to have lost lethality, this implies that a mutation has occurred in another location in the genome to prevent the effect of CcdB.

**Figure 6.**
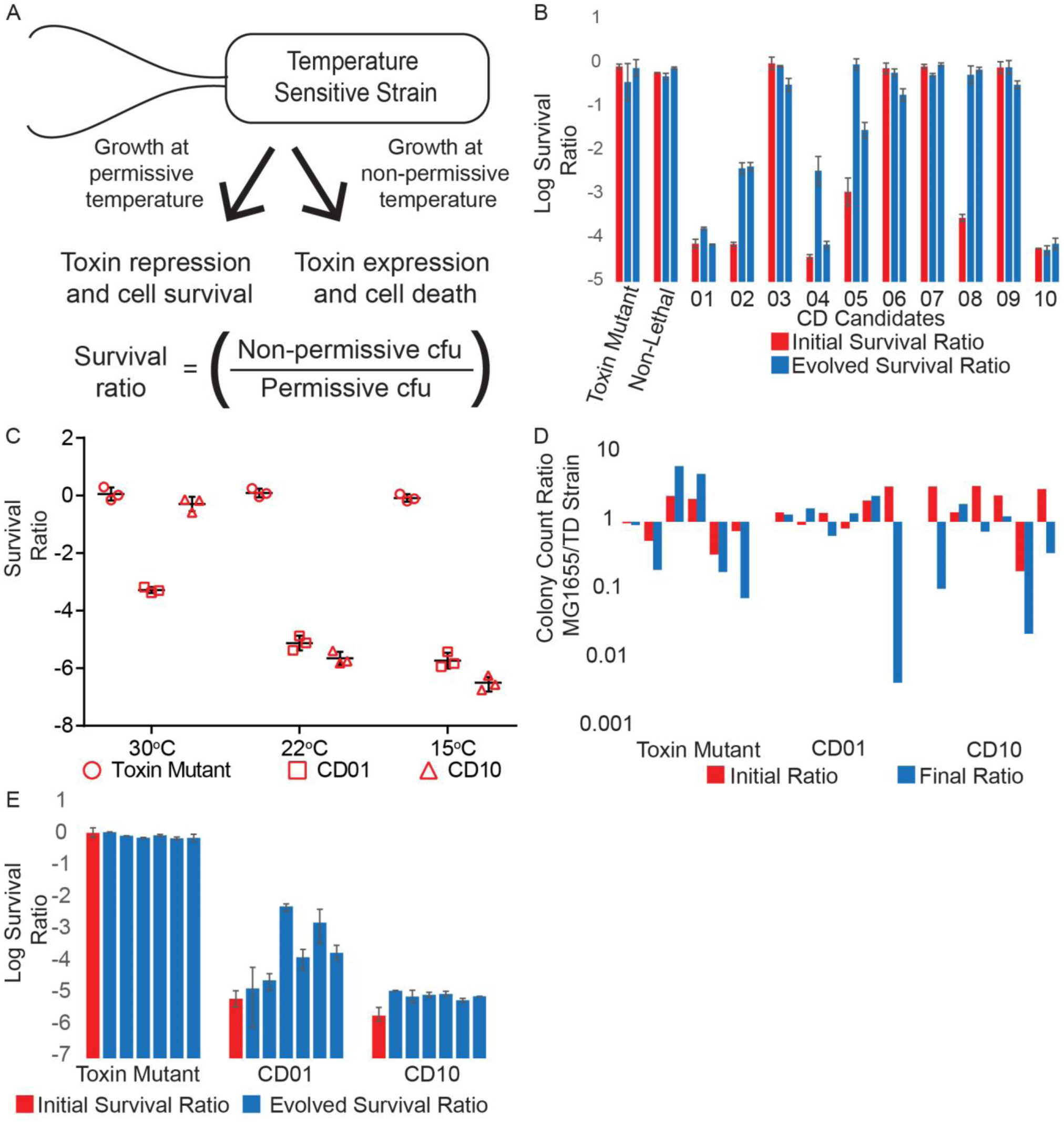
Analysis of cryodeath cadidates. **A)** Outline of the survival assay used to test the extent of population termination at non-permissive temperatures. **B)** Survival ratio of the ten temperature sensitive candidates in DH10β (CD1-CD10) before and after 140 generations of growth at a permissive temperature. Data represents average of two technical repeats, with error bars showing range. Two biological repeats after a period of growth are shown. Non-Lethal = CD non lethal. **C)** Survival ratio of candidates CD01 and CD10 in MG1655 at 30 °C, 22 °C, and 15 °C. Three technical repeats at each condition. **D)** Six biological repeats of a competitive growth assay to compare the fitness of parental (MG1655) and engineered bacterial strains. Mixed cultures were titered before and after sub-culturing for 70 generations. **E)** Six biological replicates of survival ratio of candidates CD01 and CD10 in MG1655 after 140 generations, each biological replicate data point is the average of 3 technical replicates plotted with 3 technical repeats. Initial survival ratios are from the data collected for figure 6C.

The cryodeath kill switch candidates were transferred to a different genetic background for use in the mammalian gut. After transfer by P1 transduction from DH10β into MG1655, a survival assay was used to measure lethality in both strains at 30 °C, 22 °C and 15 °C. In DH10β, both CD01 and CD10 induced no death compared to the toxin mutant control at 30 °C, a survival ratio ranging from 10^-4^ to 10^-5^ at room temperature and 10^-5^ at 15 °C (Supplementary Figure 3). In MG1655, survival ratios were over an order of magnitude lower, 10^-5^ to 10^-6^ at 22 °C, and 10^-7^ for candidate CD10 at 15 °C (Figure 6C). Candidate CD01 also had a substantial drop in survival ratio at 30 °C, (10^-3^).

The cryodeath candidates showed no disadvantage in growth rate when compared to wild type MG1655. Co-cultures were grown in minimal media for approximately 70 generations. The population ratio of kill switch strain to MG1655 was measured by comparing cfu before and after growth across 6 biological repeats (Figure 6D). Across the repeats, it varied which strain gained an evolutionary advantage over the time period, implying that a spontaneous mutation independent of the kill switch was responsible for the advantage.

One cryodeath strain was stable after an extensive period of growth. An evolutionary stability experiment was repeated with 6 biological repeats of CD01, CD10 and the toxin mutant control. Again the candidates were grown for approximately 140 generations and then assayed at room temperature to determine their evolved survival ratio. CD10 maintained its survival ratio at around 10^-5^ after the period of growth across all 6 biological repeats. The survival ratio for CD01 decreased by at least an order of magnitude in four cultures (Figure 6E) suggesting that a certain proportion of the population had lost or modified the lethal phenotype. As the candidate displaying the most robust evolutionary stability, CD10 was chosen as the most desirable cryodeath construct and was tested in the mammalian gut.

CD10 induced efficient and stable population death upon defecation from the mammalian gut. Streptomycin resistance was introduced to CD10 and the toxin mutant control in order to facilitate colonization of the mouse gut by P1 transducing a mutated *rpsL* from a spontaneously streptomycin resistant strain. Six Balb C mice, for 3 biological repeats of each construct, were treated with streptomycin throughout the experiment. CD10 and the toxin control were gavaged into separate mice after 24 hours of streptomycin treatment, and fecal samples were collected before gavaging, 24 hours after gavaging and again after 10 days. The survival assay was then carried out on cultures grown from all fecal samples, showing a similar level of lethality, a survival ratio of -5, across all mice and time points to that already described for in vitro experiments (Figure 7B). No bacteria were isolated on streptomycin plates from the sample collected before gavaging.

**Figure 7:**
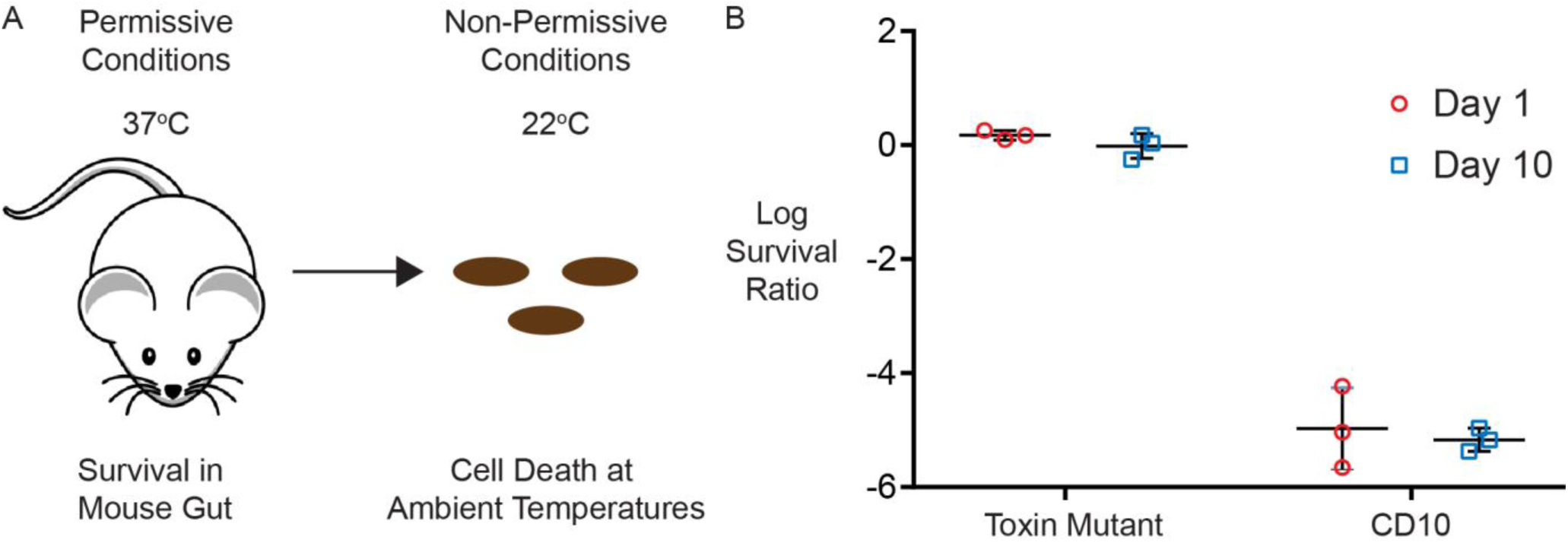
Testing cryodeath in vivo. **A)** Containment of transgenic bacteria to mammalian gut by temperature sensitive induction of cryodeath. **B)** Survival assay of cultures grown from feces of 3 mice gavaged with Toxin Mutant and three mice gavaged with CD10 in MG1655, both 1 day and 10 days after gavaging. Each data point is a separate biological repeat, made from the average of three technical repeats.

## Discussion

We have constructed two inducible toxin/antitoxin kill switch systems for controlling the environment or conditions in which a genetically engineered strain of *E. coli* can survive (Figure 1). The first is the “essentializer” element, a kill switch that links cell survival to the presence of the memory element, an engineered genetic circuit that is normally not essential (Figures 2-3). The second, “cryodeath,” responds to environmental temperature, allowing growth at 37°C but resulting in a survival ratio of less than 10^-5^ at 22°C and below (Figures 6 and 7). Using these two systems, it is possible to limit the growth conditions of transgenic *E. coli* without the need for human monitoring or input.

The design criteria for these kill switches involve multiple phenotypes, including a lack of detrimental effect on viability and growth, which collectively do not lend themselves to high-throughput screening. Our approached therefore relied on rational design and construction of small libraries whose members could be extensively tested. A novel technique for varying promoter and ribosome binding site expression levels was used to create a pool of candidates with different levels of toxin and antitoxin production. Overlap extension PCR or Gibson assembly with degenerate primers were used to vary primarily those bases expected to modulate the quantitative level of gene expression but not affect regulation. This method was chosen over random mutagenesis technique such as error prone PCR, as the limited pool of potential candidates could be screened in its entirety if needed, and because an intense random mutagenesis would likely create primarily loss-of-function mutations that could mask mutations of interest. In addition, as many of the possible nucleotides have a well characterized effect on expression (Mulligan et al., 1984; R K Shultzaberger et al., 2001; Ryan K. Shultzaberger et al., 2007), we could limit the potential candidates to those that were more likely to perform within the desired range of expression. Another alternative approach, use of wild-type promoters with varying strengths, was not chosen because such promoters may be regulated in an unknown manner, and because the chosen approach was technically simpler, faster, and less expensive. The success of modifying two different regulatory regions indicates this technique could be applied to any regulatory region where the influence of each individual base can be identified with a reasonable degree of accuracy, i.e. any promoter in *E. coli* or other well characterized organisms. Due to the error prone nature of oligo synthesis, many of the constructs sequenced had mutations likely to affect regulatory function, including the most stable cryodeath candidate CD10.

Both kill switches maintain functionality after at least 140 generations (20 passages) *in vitro* (Figure 3 and 6). The cryodeath CD10 element maintained 100% of its functionality over this time (Figure 6). In addition, CD10 maintained its lethality when used *in vivo* to limit bacterial growth after expulsion from the mammalian digestive tract in a mouse model. Under these conditions, cryodeath remained functional for at least 10 days (Figure 4), approximately 80 generations (Myhrvold, Kotula, Hicks, Conway, & Silver, 2015). Of the 10 cryodeath strains that originally showed signs of lethality, only 1 proved to be stable after an opportunity to evolve as opposed to 10 out of 11 essentializer candidates. This can likely be attributed to two causes; a lower basal expression level from P_*R*_ compared to P_*cspA*_, and the greater opportunity for the unstable essentializer candidates to lose their lethality throughout the more extensive screening process.

Our system consistently reported a survival ratio of at least 10^-5^ at 22°C (Figure 3 and 4) and can reach as low a survival ratio as 10^-6^ at 15^°^C (Figure 3C). P_*cspA*_ has been shown to induce a 16 fold increase in expression as high as 27°C, indicating the potential for a lethal level of induction at higher temperatures (Hoynes-O’Connor, Shopera, Hinman, Creamer, & Moon, 2017). Previously published kill switches using the toxin CcdB as the sole source of cell death in *E. coli* have a survival ratio of approximately 10^-3^ (Chan et al., 2015; Piraner et al., 2016). This disparity in level of lethality can be partially explained by the fact that the target of CcdB, the GyrA subunit of DNA gyrase, is one of the genes upregulated by the cold shock response (Jones, Krah, Tafuri, & Wolffe, 1992). In addition, the fact that our approach yielded a kill switch that induced a higher rate of population death can be attributed to the screening of a rationally designed small library.

An important design requirement is that kill switches should not be deleterious to growth of the host organism in permissive conditions. This means that stochastic variation in expression of the toxin and antitoxin genes should not allow greater expression of toxin even in rare circumstances. Natural toxin-antitoxin systems avoid spontaneous deletion because the antitoxin is much less stable than the toxin, so a stochastic decrease in antitoxin expression will not be significantly averaged over time. The antitoxin promoter used here is based on the *lac* promoter, which initiates transcription about once per minute when induced (Young & Bremer 1975), corresponding to an average of 20 events/generation in rapidly growing cells. A cell will have a 2-fold reduction in antitoxin expression in about 1% of all such periods (P(< =10) when the average is 20), so that if the toxin is regulated by a promoter with an induction ratio of >5, our mutation-and-screening approach should yield satisfactory kill-switches with a high probability.

A variety of promoters responding to different environmental conditions have been identified in *E. coli*, and our system could be modified to include an alternative mode of death induced by pH (Chou, Aristidou, Meng, Bennett, & San, 1995), oxygen level (The, 1990), or nutrient abundance (Yansura & Henner, 1990). Combined with an independent toxin-antitoxin system (Yamaguchi & Inouye, 2011), the lethality of the system would be increased whilst maintaining a high level of evolutionary stability (Torres, Jaenecke, Timmis, García, & Díaz, 2003).

## Contributions

Conceptualization, F.S. and J.W.; Investigation, F.S., L.B. and J.O.; Writing – Original Draft, F.S.; Writing – Review & Editing, F.S., J.W., P.S., L.B., and J.O.; Funding Acquisition, P.S. and J.W.

## Acknowledgements

This work was supported by Defense Advanced Research Projects Agency Grant HR0011-15-C-0094 and funds from the Wyss Institute for Biologically Inspired Engineering.

## Methods

### Bacterial strains

The K12 strain MG1655 was used as the bases for all strains, except for the initial screen of cryodeath constructs that was conducted in DH10β. The recombinant strain TB10 is a derivative of MG1655, with a large section of the lambda prophage genome inserted into biotin operon. In addition, the lambda *red* genes α, β and γ are under the control of cI^857^, making them temperature inducible. PAS132 contains the previously published memory element (Kotula et al., 2014) and accompanying trigger element.

### Media and growth conditions

Unless otherwise specified LB media with relevant antibiotic was use for all growth conditions. For the competitive growth assay, screening for lethality of the essentializer candidates and for the initial evolutionary screen of the essentializer candidates, M9 minimal media supplemented with 1 mM MgSO_4_, 1 μg/ml thiamine hydrochloride, 0.4% w/v glucose, 100 μMCaCl_2_ was used. For all evolution experiments and survival assays no antibiotic were used (except for streptomycin in the fecal sample survival assay). For mouse survival assays, MacConkey lactose with streptomycin was used. For plating the competitive growth assay, MacConkey rhamnose without antibiotic was used to differentiate between wild type MG1655 and strains with kill switches inserted into the rhamnose operon. Unless otherwise stated, in all instances of antibiotic use the following concentrations were employed: Amp 100 μg/ml, Kan 50 μg/ml, Gent 10 μg/ml, Strep 100ug/ml.

### Cold-sensitive reporter plasmid design

The P_*cspA*_ regulatory region was amplified using PCR from the K12 genome using primers TS3 and TS4 or TS6 to add homology to the plasmid pUA66, and incorporate the GGGGS linker in the case of TS6. pUA66 GFP was linearized by PCR using primers TS1 and TS2 and combined with P_*cspA*_ promoters to create plasmids with temperature sensitive expression of GFP.

### Construction of kill switch libraries

To construct essentializer variants (figure 2C) a cassette was ordered as a Gblock from Integrated DNA Technologies with the sequence “original EE sequence” (supplementary table 1). The degenerate oligo FS1 as well as primers FS2, FS3 and FS4 were used to amplify the cassette in two parts, which were subsequently combined using overlap extension PCR and amplified with FS3 and FS4. For the cryodeath kill switch, the degenerate primers TS12 and TS13 or TS18 were used to amplify P_*cspA*_ with and without a linker respectively. Primers TS14-TS17 were used to amplify “original EE sequence” in two parts, which were subsequently combined with the degenerate P_*cspA*_ amplicons using Gibson assembly and amplified using TS14 and TS17. In initial experiments, we found that the stitched product appeared to be correct based on gel electrophoresis, but failed to produce transformants upon recombineering (see below). We hypothesized that during the stitching amplification, the amount of DNA product exceeded the dNTPs and/or primers, with the result that the final PCR cycles simply involved denaturation and reannealing of full-length single strands, which would in general contain numerous mismatches. Upon transformation and recombination into the genome, such mismatches would be expected to undergo mismatch repair in a strand-independent manner, resulting in double-stranded breaks as gap-repairing polymerases meet. To avoid this problem, the reaction was passed through a Zymo Clean and Concentrator^TM^ kit before being added to fresh PCR reagents for an additional cycle with primers FS3 and FS4 or TS14 and TS17 for the essentializer and cryodeath libraries respectively. This ensured that the predominant PCR product was a matching double stranded helix. After this modification, the frequency of transformant isolation increased dramatically.

### P1 transduction

P1 lysates conducted according to previously published methods (L. C. Thomason, Costantino, & Court, 2007) with lysate of greater than 10^9^ pfu.

### Recombineering

For both kill switches, electrocompetent cells were prepared for transformation using previously published methds (L. Thomason et al., 2007). The degenerate essentializer library was transformed into the recombinant strain TB10 that had been induced for 15 minutes in a 42°C water bath with manual shaking every two minutes. For the cryodeath degenerate library, the recombinant plasmid pKD46 (Datsenko & Wanner, 2000) was transformed into Thermo Scientific MAX Efficiency DH10β electrocompetent cells. Cells were induced for 2-3 hours in the presence of 10 mM L-arabinose. Approximately 100 ng of DNA was combined with 50 μl of cells in a 0.1 cm cuvette, and electroporated using EC1 setting on a Biorad Micropulser. Cells were recovered in SOC medium for one hour before being spread on the relevant antibiotic plate.

### Temperature sensitive survival assay

All colonies from the recombination of the two temperature sensitive libraries (with and without a linker) were grown at 37°C, 30°C, 22°C and 15°C. 5ml overnight cultures were made at 37°C of all samples from the 37°C plate, and combined 1:1 with 50% glycerol at 37°C before being immediately transferred to a -80°C freezer. Unless being assayed for growth at lower temperatures, all subsequent experiments involving temperature sensitive strains were prepared in a constant 37°C environment. For the strains that showed variation of growth at different temperatures, the glycerol stocks were used to make overnight cultures at 37°C in 5ml of LB. This was diluted by a factor of 10 into 4 ml of LB in a 15 ml culture tube, and grown at 37^°^C until ∼OD_600_ 1.0 was reached. Serial dilutions of the cultures were made and plated onto a number of LB plates corresponding to the number of temperatures being assayed. Separate plates were then incubated for up to 15 days at various temperatures, and the cfu at each temperature compared to the cfu at 37°C was termed the survival ratio.

### Testing evolutionary stability of kill switch strains

For the essentializer candidates, glycerol stocks were made of the 11 candidates that proved lethal from extensive screening outlined in figure 2E. These glycerol stocks were used to inoculate 2 ml of M9 minimal media without antibiotic. After 24 hours of growth, 20 μl was used to inoculate 2 ml of M9 minimal media. This was repeated for 20 passages. The assay in figure 3A was then used to determine whether the essentializer was still producing a selective pressure to keep the memory element. For the cryodeath candidates, the glycerol stocks of the temperature sensitive candidates was used to inoculate 2 ml of LB without antibiotic. After growth over night, 20 μl was used to inoculate 2 ml of LB, which was grown for 4 hours twice by diluting the same way. This cycle of overnight, 4 hours of growth, 4 hours of growth was repeated 7 times. Glycerol stocks of the final cultures were made, and used to inoculate cultures for the survival assay.

### Competitive Growth Assay

Overnight cultures of MG1655, Toxin Mutant, EE10, EE11, CD01 and CD10 were grown to stationary phase. 100ul of each culture was then used to inoculate M9 minimal media, and glycerol stocks were made of this initial inoculation. Cultures were passaged once every 24 hours diluting 1:1000 into M9 minimal media. After 8 days, glycerol stocks were made of the final culture. Glycerol stocks were then defrosted and plated directly onto MacConkey Rhamnose plates. As the two kill switches had been inserted into the Rhamnose operon, they could be distinguished as making white colonies on Rhamnose MacConkey compared to red colonies produced by MG1655. Cfu were used to determine the ratio of kill switch strain to MG1655.

### Testing Temperature Sensitive Kill Switch in the Mammalian Gut

Approval for animal work came from the Harvard Medical School IACUC under protocol 04966. Seven week old female BalbC mice from Charles River Laboratories were given two weeks to acclimatize to the facility upon delivery. 500 ug/ml of streptomycin was added to their drinking water and maintained throughout the experiment. The Toxin mutant and strain CD10 were gavaged at a concentration of approximately 10^8^ cfu/ml, each into 3 separate mice. Feces was collected between 2-6pm from day 0 (immediately before gavaging) to day 10, and placed on dry ice before being transferred to -80 freezer. Feces was resupended in PBS by shaking at 4 °C for 1 hr, then 100 μl was used to inoculate 5ml of MacConkey lactose broth with streptomycin and grown overnight. The following day this overnight culture was diluted by a factor of 10 into MacConkey lactose and grown to approximately OD 1.0. Cultures were then serial diluted and plated at 37 °C and room temperature to compare cfu at permissive and non-permissive temperatures.

## Supplementary figures and tables legends

**Supplementary figure 1.**
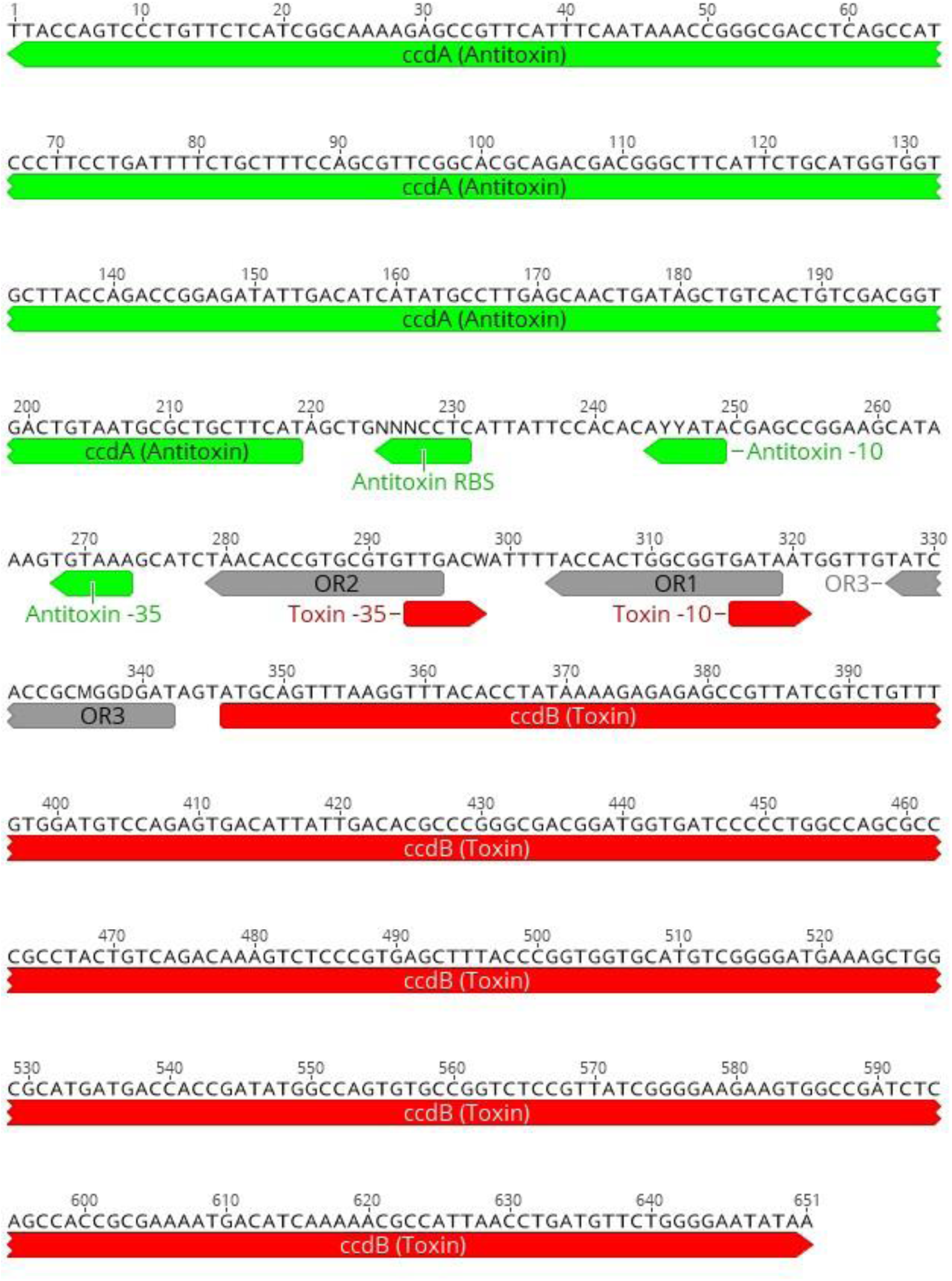
Related to figure 2: Sequences of essentializer degenerate library. Green = the antitoxin ORF and regulatory regions, Red = the toxin ORF and regulatory regions. Grey = the binding regions for cI and Cro, * indicating it has been modified to increase binding efficiency according to previously published work (Sarai & Takeda, 1989; Takeda et al., 1989).

**Supplementary table 1.**
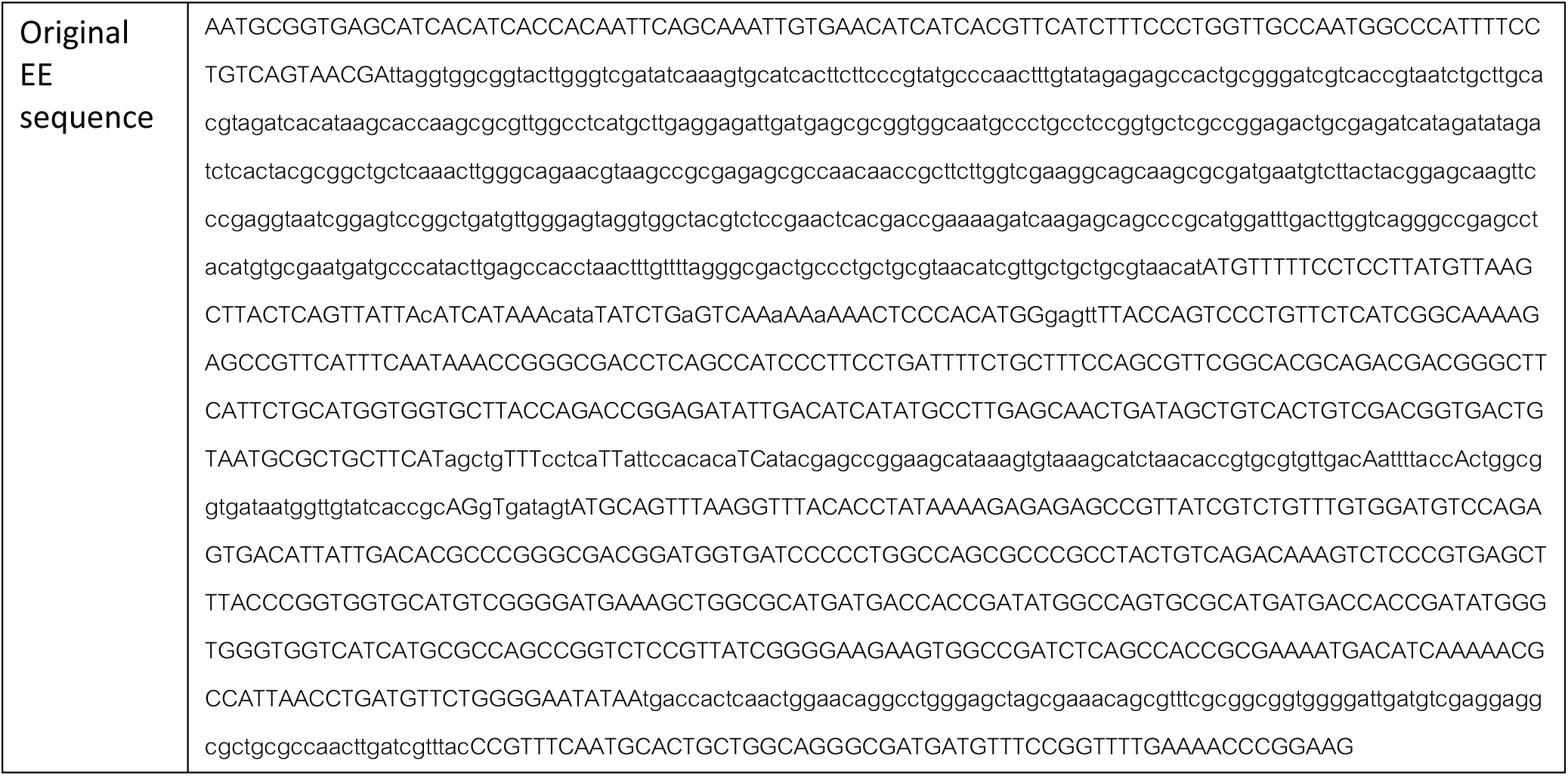
Related to figure 2: Sequence ordered as a G block from Integrated DNA technologies. Includes the sequence displayed in supplementary figure 1 as well as a gentamycin resistance cassette and 100 bp of homology on either end to the rhamnose operon.

**Supplementary table 2.** Related to figure 2 and 3: Nucleotide located at each degenerate base location for EE01-EE11, going from 5’-3’. *indicates when a deletion event resulted in this base in this approximate position.

|  | N | N | N | Y | Y | W | M | D |
| --- | --- | --- | --- | --- | --- | --- | --- | --- |
| EE01 | T | G | A | T | T | T | A | A |
| EE02 | G | A | A | A* | A* | T | A | G |
| EE03 | A | G | G | C | T | T | A | A |
| EE04 | C | G | C | C | T | T | A | A |
| EE05 | T | C | G | T | T | T | A | A |
| EE06 | G | A | A | C | T | T | C | A |
| EE07 | G | C | G | C | C | A | C | A |
| EE08 | G | G | A | C | T | A | C | A |
| EE09 | G | T | G | T | C | A | A | G |
| EE10 | A | T | G | T | T | A | C | A |
| EE11 | C | G | T | C | T | T | C | A |

**Supplementary table 3.**
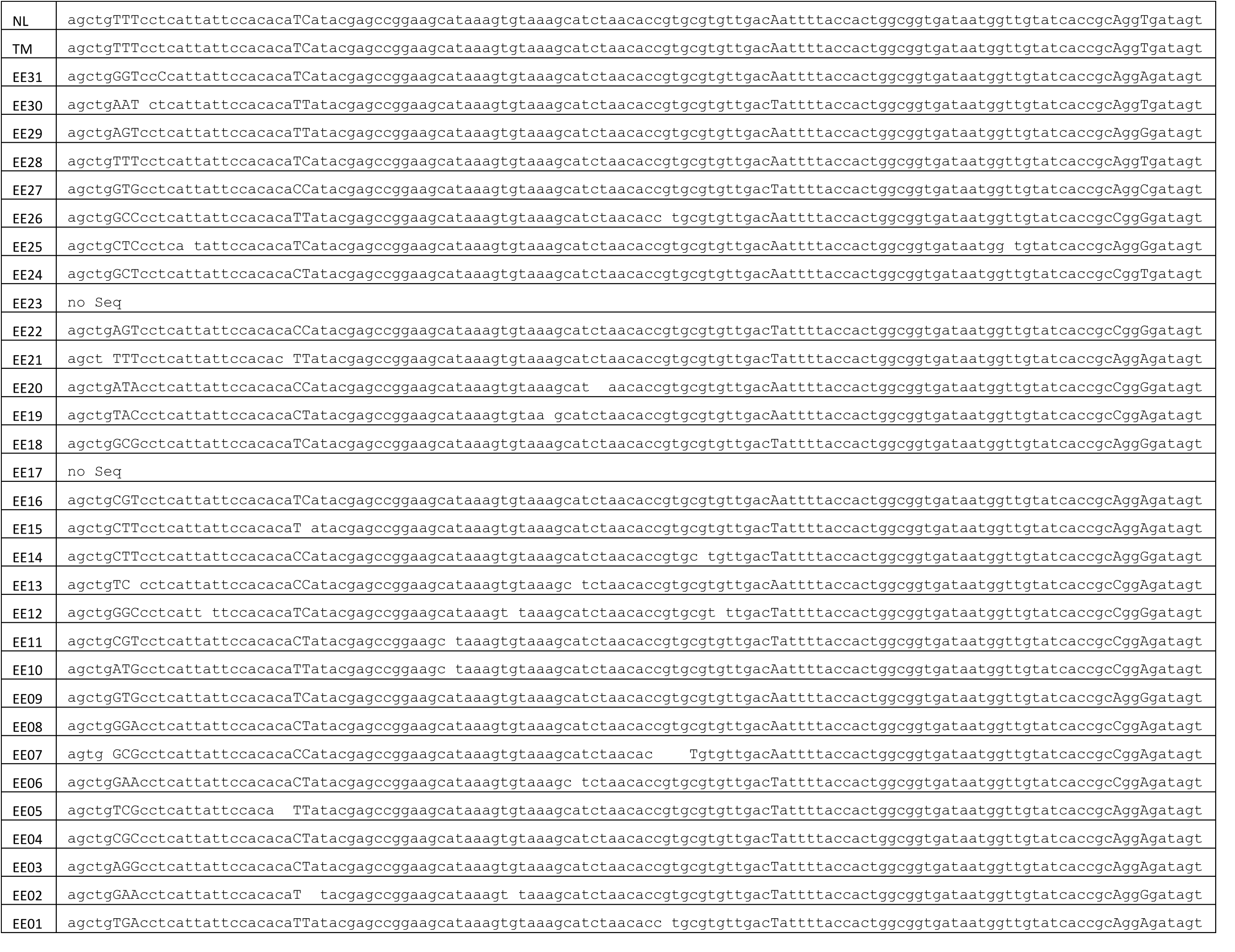
Related to figure 2 and 3: complete sequence of regulatory region between the start codon of the toxin and the start codon of the antitoxin for all essentializer candidates analyzed. NL = non lethal, TM = toxin mutant, EE01-EE11 = lethal candidates. EE12-EE40 = non-lethal candidates.

**Supplementary table 4.**
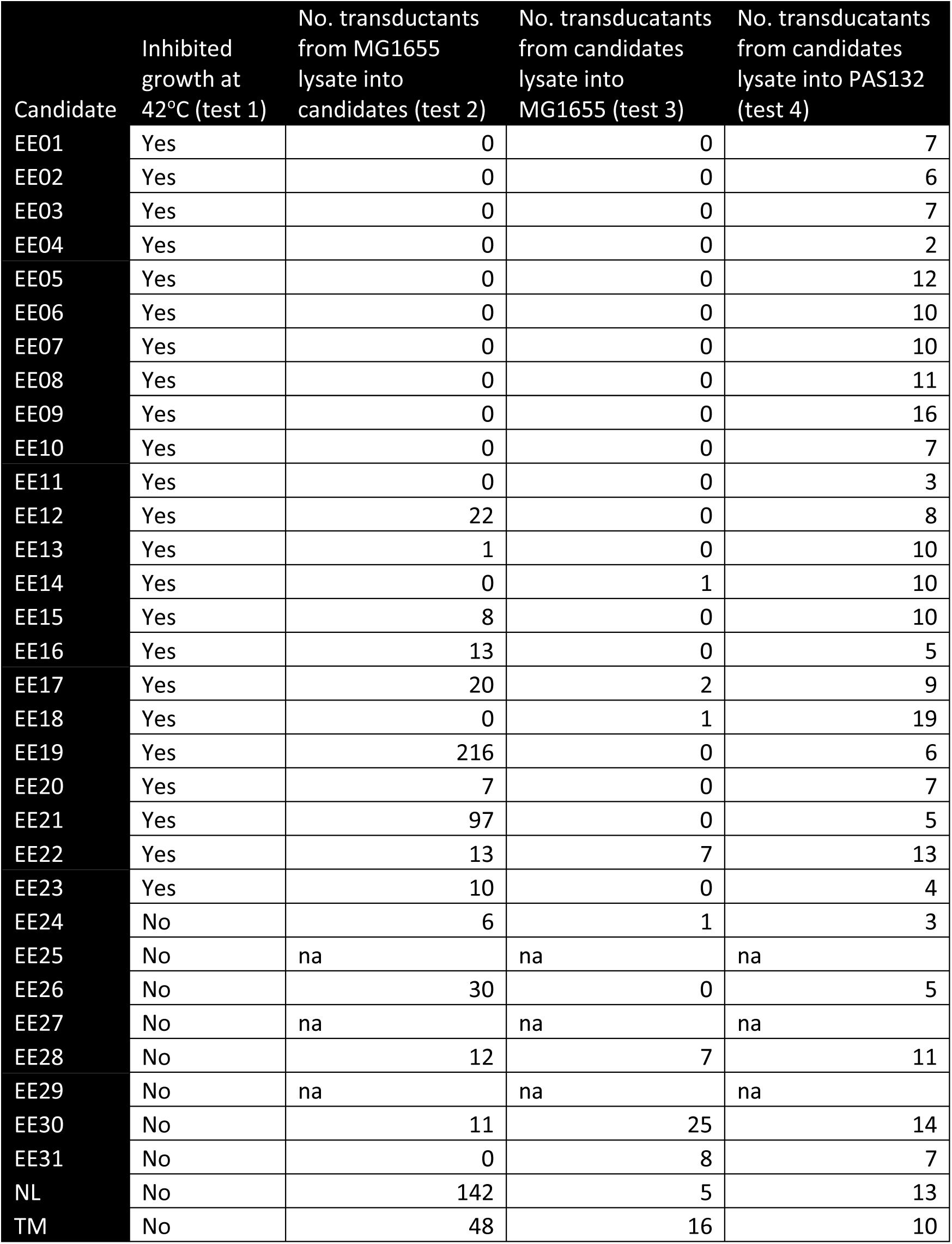
Related to figure 2: Raw data from tests 1-4 of flowchart in figure 2E.

| Candidate | Inhibited growth at 42°C (test 1) | No. transductants from MG1655 lysate into candidates (test 2) | No. transducants from candidates lysate into MG1655 (test 3) | No. transducants from candidates lysate into PAS132 (test 4) |
| --- | --- | --- | --- | --- |
| EE01 | Yes | 0 | 0 | 7 |
| EE02 | Yes | 0 | 0 | 6 |
| EE03 | Yes | 0 | 0 | 7 |
| EE04 | Yes | 0 | 0 | 2 |
| EE05 | Yes | 0 | 0 | 12 |
| EE06 | Yes | 0 | 0 | 10 |
| EE07 | Yes | 0 | 0 | 10 |
| EE08 | Yes | 0 | 0 | 11 |
| EE09 | Yes | 0 | 0 | 16 |
| EE10 | Yes | 0 | 0 | 7 |
| EE11 | Yes | 0 | 0 | 3 |
| EE12 | Yes | 22 | 0 | 8 |
| EE13 | Yes | 1 | 0 | 10 |
| EE14 | Yes | 0 | 1 | 10 |
| EE15 | Yes | 8 | 0 | 10 |
| EE16 | Yes | 13 | 0 | 5 |
| EE17 | Yes | 20 | 2 | 9 |
| EE18 | Yes | 0 | 1 | 19 |
| EE19 | Yes | 216 | 0 | 6 |
| EE20 | Yes | 7 | 0 | 7 |
| EE21 | Yes | 97 | 0 | 5 |
| EE22 | Yes | 13 | 7 | 13 |
| EE23 | Yes | 10 | 0 | 4 |
| EE24 | No | 6 | 1 | 3 |
| EE25 | No | na | na | na |
| EE26 | No | 30 | 0 | 5 |
| EE27 | No | na | na | na |
| EE28 | No | 12 | 7 | 11 |
| EE29 | No | na | na | na |
| EE30 | No | 11 | 25 | 14 |
| EE31 | No | 0 | 8 | 7 |
| NL | No | 142 | 5 | 13 |
| TM | No | 48 | 16 | 10 |

**Supplementary figure 2.**
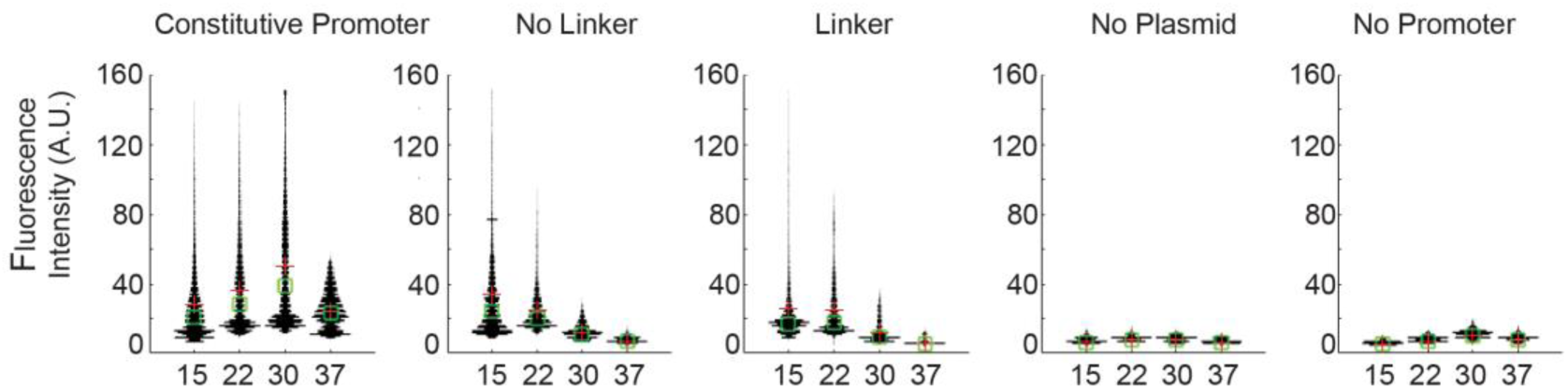
Related to figure 4: Fluorescence intensity for images in figure 4B. Images were partitioned into 5 section between streaked bacteria. A histogram of fluorescence intensity using arbitrary units is plotted for the strains containing different plasmids at a range of temperatures.

**Supplementary figure 3.**
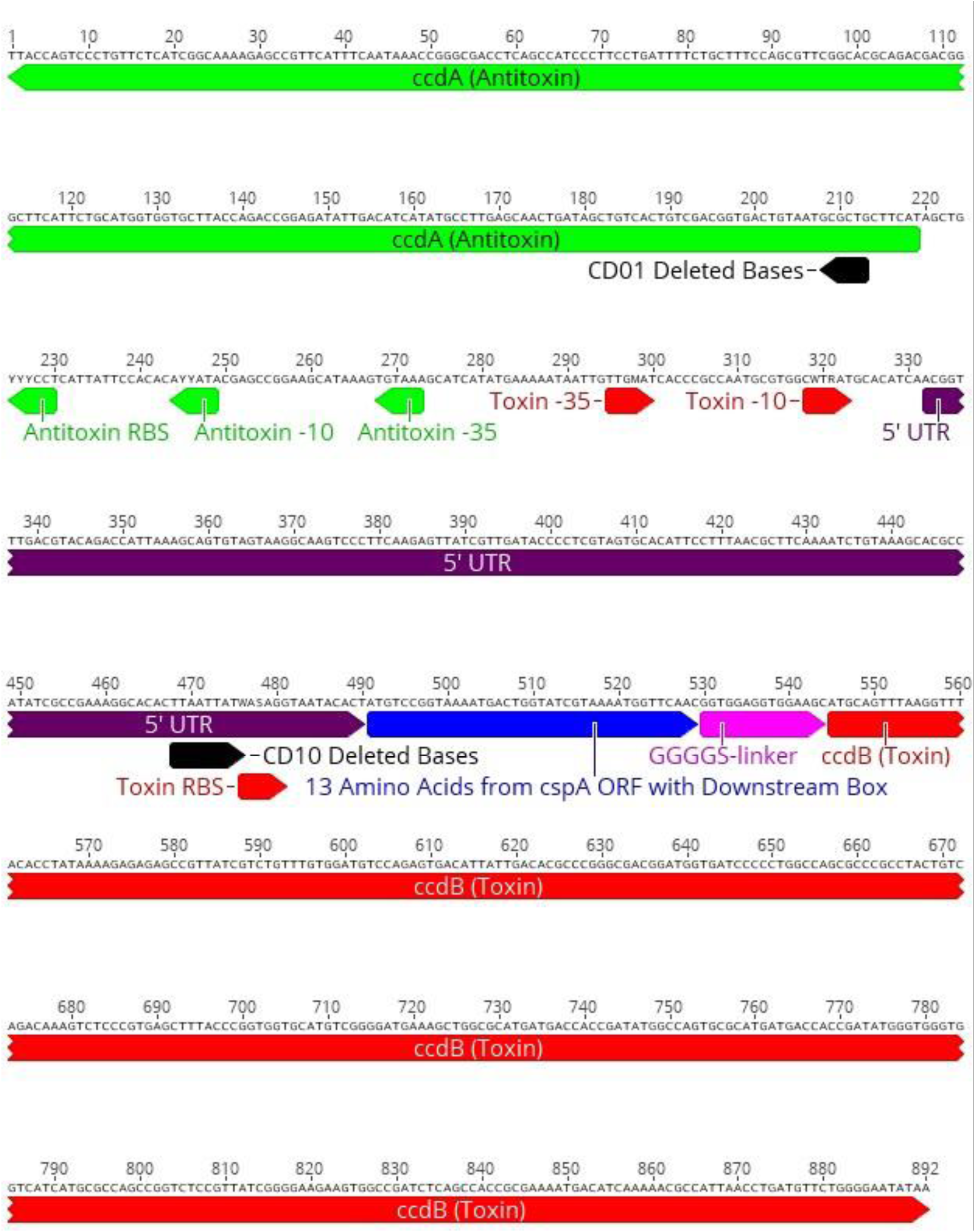
Related to figure 5: Sequence of cryodeath degenerate library. Green = the antitoxin ORF and regulatory regions, Red = the toxin ORF and regulatory regions. Purple = toxin long 5’ UTR. Blue = first 13 amino acids of *cspA* ORF, including the downstream box. Purple is the GGGGS linker that is present in all candidates except CD1 and CD11. Black annotations represent the regions that are deleted in CD01 and CD10.

**Supplementary table 5.** Related to figure 5 and 6: Nucleotide located at each degenerate base location for CD01-CD10, going from 5’-3’. *indicates when a deletion event resulted in this base in this approximate position.

|  | Y | Y | Y | Y | Y | M | W | R | W | S |
| --- | --- | --- | --- | --- | --- | --- | --- | --- | --- | --- |
| CD01 | T | C | T | T | C | C | T | G | A | C |
| CD02 | C | T | T | C | T | A | T | G | T | C |
| CD03 | T | T | T | T | C | C | T | G | T | C |
| CD04 | C | T | T | C | T | A | T | G | T | C |
| CD05 | T | C | C | T | T | C | T | G | A | C |
| CD06 | T | T | T | T | C | A | T | G | T | C |
| CD07 | T | T | T | T | C | A | T | G | A | C |
| CD08 | T | T | T | C | T | A | T | G | T | C |
| CD09 | T | T | T | C | T | A | T | G | A | C |
| CD10 | T | C | C | T | C | C | T | G | C* | G |

**Supplementary table 6.**
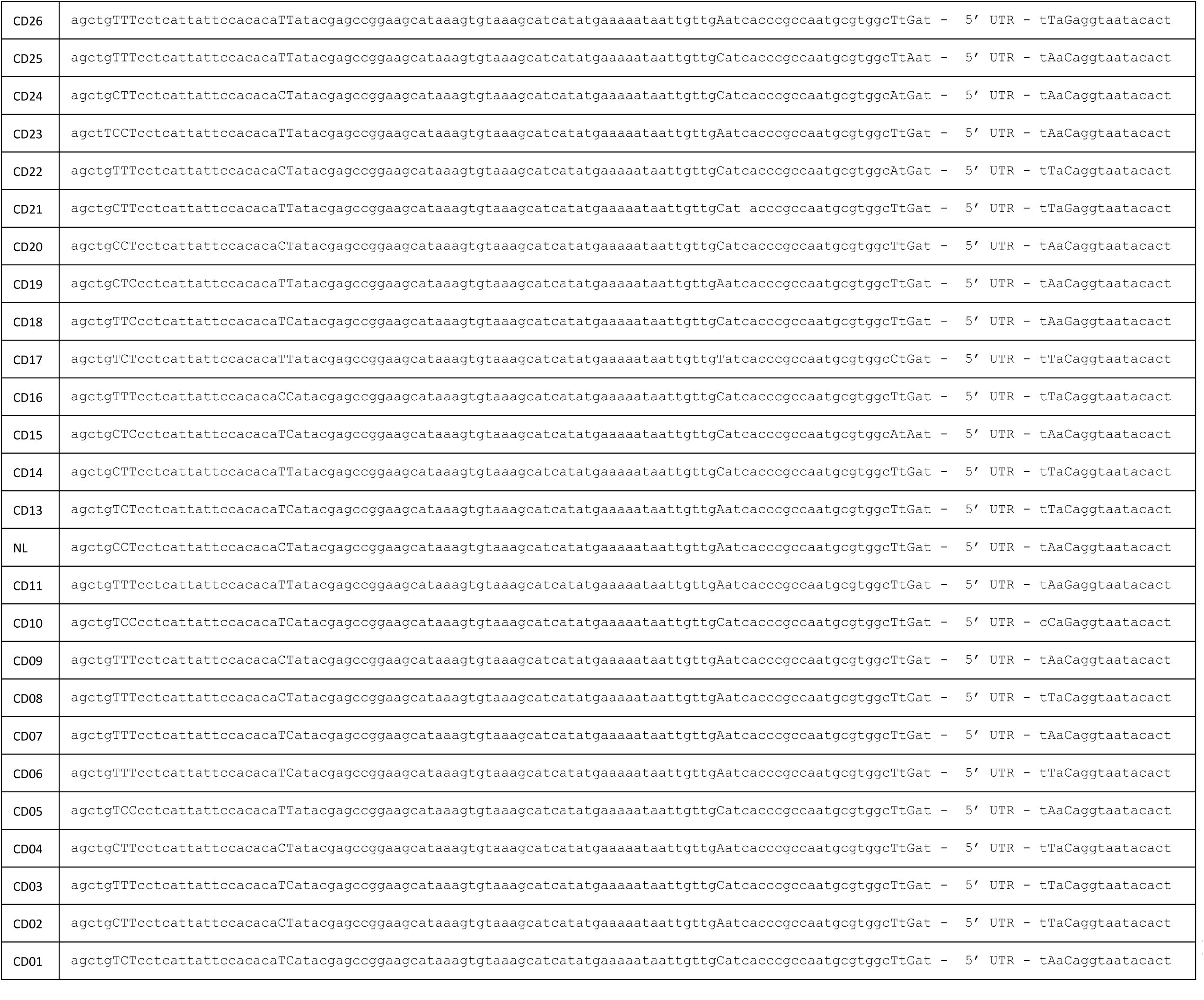
Related to figure 5 and 6: complete sequence of regulatory region minus the 5’ UTR between the start codon of the toxin and the start codon of the antitoxin for all cryodeath candidates analyzed. NL = candidate used as non-lethal control, CD01-CD10 = lethal candidates. CD11-CD26 = non-lethal candidates. Capitalized bases are were degeneracy was introduced. Sequence of 5’ UTR can be found in Supplementary table 7.

**Supplementary table 7.**
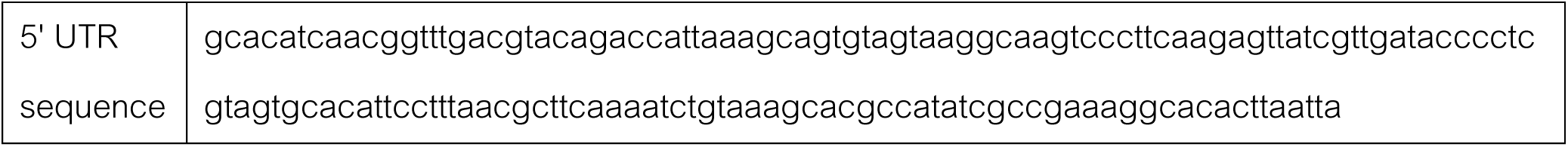
Related to figure S6: sequence of 5’ UTR. Although technically not part of the 5’ UTR, bases -1 to -8 of the P_*cspA*_ are also grouped into this sequence for space considerations in supplementary figure 6.

**Supplementary Figure 4.**
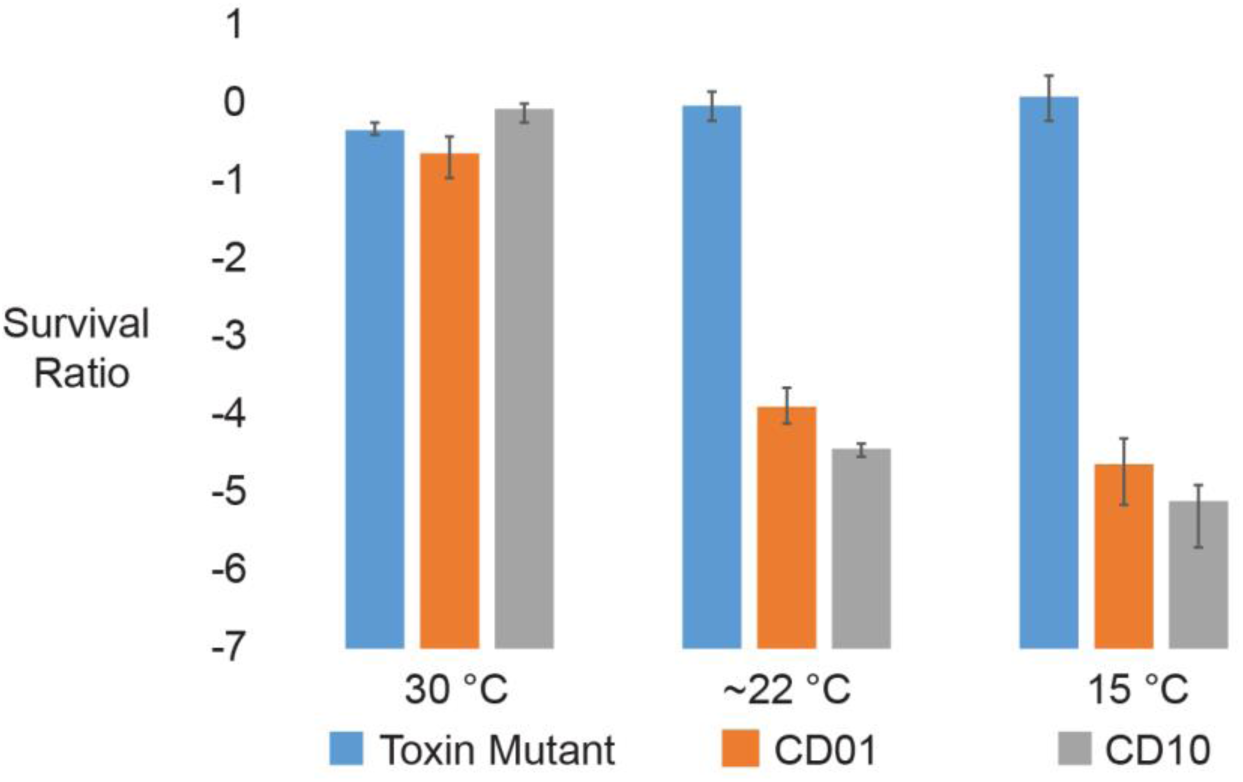
Related to Figure 6: Survival ratio of candidates CD01 and CD10 in Dh10β at 30 °C, room temperature (∼22 °C), and 15 °C. Data points are from average of 3 technical repeats.

**Supplementary table 8.**
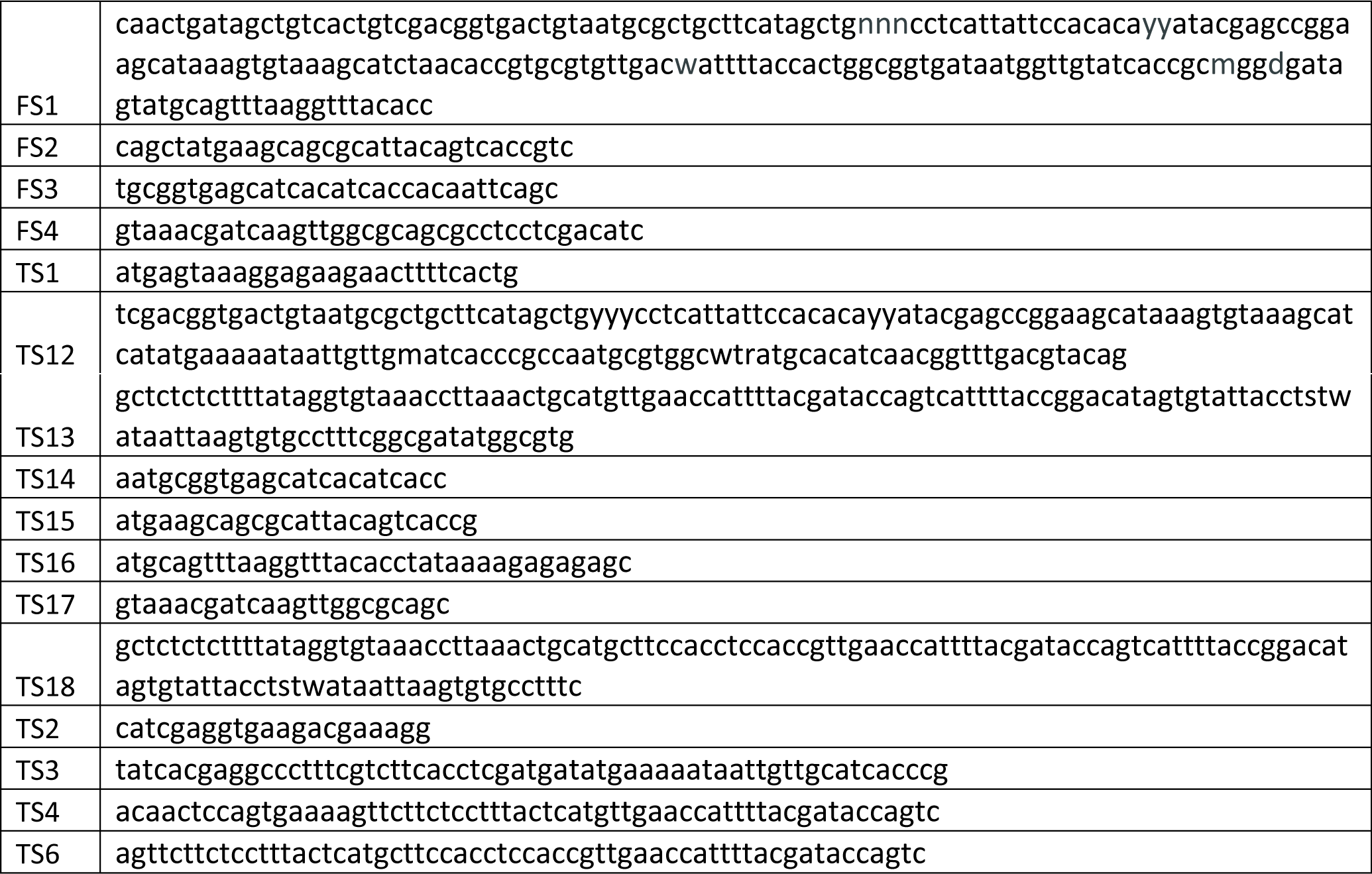
Related to Methods: Full sequence of all oligos described in methods.

